# Generation of a new *Adar1p150^-/-^* mouse demonstrates isoform-specific roles in embryonic development and adult homeostasis

**DOI:** 10.1101/2022.08.31.506069

**Authors:** Zhen Liang, Carl R Walkley, Jacki E Heraud-Farlow

## Abstract

The RNA editing enzyme Adenosine deaminase acting on RNA 1 (ADAR1) is an essential regulator of innate immune activation by both cellular and viral dsRNA. Adenosine-to-Inosine (A-to-I) editing by ADAR1 modifies the sequence and structure of endogenous dsRNA and masks it from the cytoplasmic dsRNA sensor melanoma differentiation-associated protein 5 (MDA5), preventing innate immune activation. Loss of function mutations in *ADAR* are associated with rare autoinflammatory disorders including Aicardi-Goutières Syndrome (AGS), defined by a constitutive systemic upregulation of type I interferon (IFN). The murine *Adar* gene encodes two protein isoforms with distinct functions: ADAR1p110 is constitutively expressed and localizes to the nucleus, whereas ADAR1p150 is primarily cytoplasmic and is inducible by IFN. Recent studies have demonstrated the critical requirement for ADAR1p150 to suppress innate immune activation by self dsRNAs, however, detailed *in vivo* characterization of the role of ADAR1p150 during development and in adult mice is lacking. We developed a new ADAR1p150-specific knockout mouse mutant based on a single nucleotide deletion that resulted in the loss of the ADARp150 protein without affecting ADAR1p110 expression. The *Adar1p150^-/-^* died embryonically at E11.5-E12.5 due to cell death in the fetal liver accompanied by an activated IFN response. Somatic loss of ADAR1p150 in adults was lethal and caused rapid hematopoietic failure, demonstrating an ongoing requirement for ADAR1p150 *in vivo*. The generation and characterization of this mouse model demonstrates the essential role of ADAR1p150 *in vivo* and provides a new tool for dissecting the functional differences between ADAR1 isoforms and their physiological contributions.

## Introduction

Adenosine to inosine (A-to-I) editing is one of the most prevalent RNA modifications in mammals. Tens of thousands to millions of A-to-I editing sites have been identified in mammalian genomes, the vast majority of which lie within repetitive elements such as SINEs and LINEs, as well as a smaller number present in coding regions and other RNAs such as miRNAs (Bazak et al., 2014; Licht et al., 2019b; Pfaller et al., 2018; Picardi et al., 2015; Tan et al., 2017). Two catalytically active ADAR enzymes have been identified, ADAR1 and ADAR2. When A-to-I editing occurs in a coding region, the physiological functions of the resultant protein may be altered because inosine is usually interpreted as guanosine by the ribosome (Basilio et al., 1962; Licht et al., 2019a; Martin et al., 1985). Recoding of the transcript of the glutamate receptor subunit *GRIA2*, at a single adenosine, is the essential physiological function of ADAR2 in mice (Higuchi et al., 2000; Higuchi et al., 1993). In contrast, the primary function of ADAR1 editing is to modify the structure and immunogenicity of cellular double-stranded RNA (dsRNA) (Eisenberg and Levanon, 2018; Walkley and Li, 2017). Alterations in ADAR1 expression or function have been linked to various pathological states, from common autoimmune diseases to cancers (Jain et al., 2019; Li et al., 2022). A subset of the severe pediatric autoinflammatory disease and Type I interferonopathy, Aicardi-Goutières syndrome (AGS), is caused by inherited *ADAR* mutations (Crow and Stetson, 2022; Rice et al., 2012). In AGS, *ADAR* mutations are most often compound heterozygote, with one allele having a mutation impacting the p150 isoform and the other impacting both p150 and p110 isoforms (Rice *et al*., 2012; Rice et al., 2017). Delineating the full spectrum of ADAR1 functions is critical to understanding the role of each isoform in disease and developing new therapeutics.

The cytosolic antiviral system of cells can distinguish endogenous (“self”) nucleic acid from foreign ones (“non-self”) (Chen and Hur, 2022; Gazquez-Gutierrez et al., 2021; Schlee and Hartmann, 2016). An established function of ADAR1 is to mark endogenous dsRNA as “self” through A-to-I substitution, thus masking dsRNA from the cytosolic RNA-sensing receptor MDA5 (Liddicoat et al., 2015; Mannion et al., 2014; Pestal et al., 2015). Activation of MDA5 and its downstream effector, mitochondrial antiviral signaling protein (MAVS), initiates a cascade of transcription of interferon-stimulated genes (ISGs) and triggers type I interferon (IFN) production and signaling. *Adar1* null (*Adar1^-/-^*) animals are embryonic lethal between E11.5 and E12.5, accompanied by activated type I IFN, ISG production and cell death across different organs, most notably failed hematopoiesis in the fetal liver (FL) (Hartner et al., 2004; Hartner et al., 2009; Wang et al., 2004). Mice expressing an editing deficient ADAR1 protein (*Adar1^E861A/E861A^*) also die embryonically at E13.5 with an elevated ISG signature and cell death in FL (Liddicoat *et al*., 2015). Deleting either MDA5 or MAVS can rescue the embryonic death of *Adar1* null animals to 2-3 days post-birth (Bajad et al., 2020; Liddicoat *et al*., 2015; Mannion *et al*., 2014; Pestal *et al*., 2015), and more strikingly, can allow survival of ADAR1 editing deficient mice to adulthood (Liddicoat *et al*., 2015). *Adar1^E861A/E861A^Ifih1^-/-^* mice survive longterm (Chalk et al., 2019; Heraud-Farlow et al., 2017). These genetic studies indicate the importance of ADAR1 protein and A-to-I editing in suppressing the MDA5 sensing pathway and preventing embryonic abnormalities. Importantly, the genetic pathways are conserved in humans. Loss of ADAR1 in human cells triggers an MDA5-dependent upregulation of the Type I IFN response (Chung et al., 2018; Pestal *et al*., 2015; Pfaller *et al*., 2018). Consistent with the genetics resolved in mouse models, patients with loss-of-function mutations in *ADAR* or gain-of-function mutations in *IFIH1* (encoding MDA5) both develop AGS (Crow and Stetson, 2022).

In both humans and mice, the *ADAR* gene encodes two protein isoforms with distinct functions, ADAR1p110 and ADAR1p150. ADAR1p110 is constitutively expressed and restricted to the nucleus, where it edits RNA co-transcriptionally (Hsiao et al., 2018; Rodriguez et al., 2012). ADAR1p150 is primarily expressed in the cytoplasm and is inducible in response to stimuli such as viral infection and type I IFN (Samuel, 2011; 2019). In some organs of the developing mouse, such as the spleen and thymus, it appears to be the predominant isoform (Kim et al., 2021). ADAR1p150 has a unique Z-DNA/RNA-binding alpha domain (Zα) at its N-terminus that can bind to and interact with the alternative left-handed conformation of RNA (Z-RNA) (Herbert et al., 1995; Herbert et al., 1998; Nakahama and Kawahara, 2021). The p150 isoform has been considered the key to suppressing MDA5-mediated sensing. This is primarily evidenced by the phenotypes of ADAR1p150-specific knockout mice (*Adar1p150^-/-^*), generated by the deletion of the first exon of the gene (Ward et al., 2011c). The *Adar1p150^-/-^* mutant was embryonic lethal at E11.0-12.0, similar to the *Adar1^-/-^* that lacks both p110 and p150 isoforms (Ward *et al*., 2011c). *Adar1p150^-/-^* embryos had abnormal morphology, and cells derived from them were more susceptible to viral infection (Ward *et al*., 2011c). Concurrent deletion of MAVS could rescue the embryonic lethality of the *Adar1p150^-/-^* mice, with animals surviving to weaning (Pestal *et al*., 2015). In contrast, mice without the p110 isoform (*Adar1p110^-/-^*) have normal embryonic development but a high post-natal mortality (Kim *et al*., 2021). The p110-deficient mice had no ISG signature indicating that the retained expression of the p150 isoform was sufficient to prevent MDA5 activation. Deletion of MDA5 in the *Adar1p110^-/-^* model (*Adar1p110^-/-^Ifih1^-/-^*) failed to prevent the high level of post-natal lethality, however, expression of a single editing dead allele of ADAR1 (*Adar1p110^-/E861A^*) could rescue viability (Kim *et al*., 2021). The above studies suggest that ADAR1p150, but not ADAR1p110, is required to suppress MDA5 activation during embryonic development.

Outside of identifying the day of embryonic lethality of the *Adar1p150^-/-^* animals, however, there was no further *in vivo* characterization of this allele.(Ward *et al*., 2011c). Furthermore, the isoform specificity of the original *Adar1p150^-/-^* allele and whether the method used for gene targeting may affect the expression of the retained ADAR1p110 expression remains an unresolved question (Matthaei et al., 2011; Steinman and Wang, 2011; Ward et al., 2011a; b; Ward *et al*., 2011c). The importance of understanding the specific functions of ADAR1p150 has been highlighted in recent years with rising interest in functions of the Zα domain, particularly as this domain is a mutational hotspot in patients with AGS (Rice *et al*., 2017). Recent studies have highlighted the functional importance of this domain in repressing the type I interferon response (de Reuver et al., 2021; Guo et al., 2021; Nakahama et al., 2021).

In this study, we have identified and characterized a new *Adar1p150^-/-^* mouse allele. This allele has a single nucleotide deletion which causes a frameshift and formation of in-frame stop codons, leading to the specific loss of ADAR1p150. Critically, this point mutation does not interfere with either the constitutive or IFN-inducible promoters of ADAR1, so the expression of ADAR1p110 remained unaffected. We have characterized the *in vivo* functions of ADAR1p150 during development and adult homeostasis to allow comparison to the ADAR1 null and editing deficient alleles. The model provides a critical tool for future studies delineating the isoform-specific effects of ADAR1 and its role in disease.

## Results and Discussion

### Identification of a new Adar1p150 specific loss of function allele

The two isoforms of ADAR1 are produced by alternative use of the first exon via different promoters. ADAR1p150 is produced from a type I IFN-inducible promoter that is located upstream of exon 1A (**Fig 1A**). The initiation of translation of the p150 isoform starts at M1 within exon 1A and the entire exon 2 functions as a coding sequence for p150. In contrast, the p110 isoform is expressed from a constitutive promoter upstream of exon 1B which is spliced into exon 2, where the p110-specific initiation codon M249 (human M296) is located (George et al., 2005). ADAR1p110 can also be produced from the IFN-inducible promoter due to inefficient translation initiation at M1, leading to the initiation of translation of p110 at M249 in exon 2 (George *et al*., 2005; Sun et al., 2021; Wong et al., 2003). The IFN-inducible expression of p110 alongside p150 may contribute to optimal editing during the IFN response (Sun *et al*., 2021). The previously described ADAR1p150 knockout mouse allele was generated by substituting the IFN-inducible promoter and exon 1A with an anti-sense-orientated pPGK-Neo cassette (**Fig 1A**) (Ward *et al*., 2011c).

**Fig 1.**
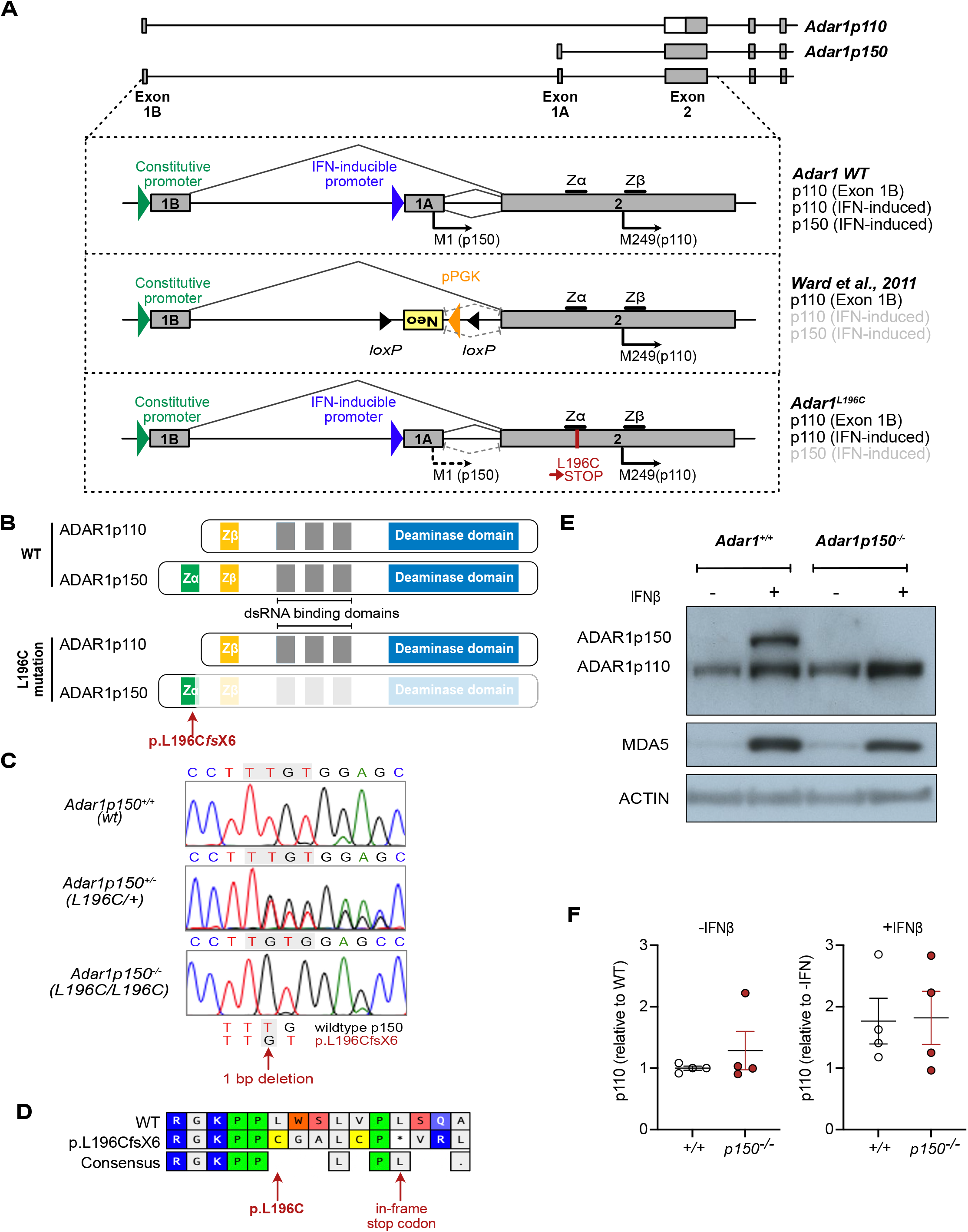
*Adar1^L196C^* mutation leads to an isoform-specific deletion of ADAR1p150 but retention of p110. **(A)** Genomic organization of the murine *Adar* locus showing locations of alternative exon 1A/1B and exon 2 and individual promoters and initiation (M1 and M249) codons that lead to expression of two ADAR1 isoforms. The IFN-inducible promoter upstream of exon 1A leads to the ADAR1p150 expression from M1, and ADAR1p110 from M249 when leaky ribosome scanning occurs. The constitutive promoter upstream of exon 1B maintains the constitutive expression of ADAR1p110. The schematic provides the comparison of the locus and protein products from wildtype *Adar*, the previous ADAR1p150-deficient model (Ward *et al*., 2011c) and the L196C allele described in this report. **(B)** Schematic of the wildtype ADAR1 isoforms p110 and p150 and the location of the L196C mutation that leads to knockout of the ADAR1p150 protein. **(C)** Sanger sequencing chromatogram and alignments of genomic DNA isolated from *Adar1^+/+^, Adar1p150^+/-^ (Adar1^L196C/+^*) and *Adar1p150^-/-^ (Adar1^L196C/L196C^*) animals indicating 1 base pair deletion in the L196C mutation. **(D)** Predicted amino acid translation of the wildtype ADAR1p150 (WT) and L196C allele. **(E)** Western blot analysis of wildtype *Adar1^+/+^* and *Adar1p150^-/-^* MEFs after 24 hours of IFN-beta (IFNβ) treatment. Expression of ADAR1 isoforms p150 and p110, MDA5 as well as ACTIN are indicated. **(F)** Quantification of the Western blots shown in Fig 1E and Supplemental Fig 1A. ADAR1p110 expression was normalized to ACTIN. The left panel shows ADAR1p110 expression at baseline normalized to *Adar1^+/+^*. The right panel shows ADAR1p110 levels plus IFN normalized to minus IFN for each cell line. N=4 for both *Adar1^+/+^* (+/+) and *Adar1p150^-/-^ (p150^-/-^*) generated from 3 independent mice. The figure presented here is the replicate of the samples in lanes 3 to 6 of Supplemental Fig1 (left image) from a different batch of IFNβ treatment. Data represent the mean +/- SEM.

During the generation of a p.P195A knock-in allele using CRISPR/Cas9 (Liang et al., 2022), we identified an incidental mutation that was predicted to result in a p150 isoform-specific knock-out allele. A single nucleotide deletion at nucleotide 587 (587delT) was identified (nomenclature based on NCBI CCDS50963). This is predicted to result in a Leucine 196 to Cysteine (p.L196C*fs*X6) amino acid change and introduction of in-frame stop codon(s) due to the change in reading frame (**Fig 1B-D**). We confirmed the introduction of the mutation in the germ-line by Sanger sequencing and then inbred heterozygous mice to assess this new allele. We generated and immortalized mouse embryonic fibroblasts (MEFs) from E11.5 embryos and treated them with murine interferon beta (IFNβ) to assess ADAR1p150 expression. The *Adar1^+/+^* MEFs expressed ADAR1p110 basally, and ADAR1p150 was robustly induced upon treatment with IFNβ. A mild elevation of ADAR1p110 protein was also observed following IFNβ treatment of *Adar1^+/+^* MEFs (**Fig 1E**). Homozygous *Adar1^L196C/L196C^* (referred to herein as *Adar1p150^-/-^* MEFs had comparable levels of ADAR1p110 to wild-type cells at baseline and had no expression of ADAR1p150 following IFNβ treatment. Importantly, the expression of ADAR1p110 was equivalent in *Adar1p150^-/-^* versus *Adar1^+/+^* MEFs, both at baseline and following induction by IFNβ treatment, indicating ADAR1p110 expression and induction was unaffected by the point mutation (**Fig 1F; Supplemental Fig 1A**). Therefore, the *Adar1^L196C^* point mutation generated a *Adar1p150* null allele without any apparent effect on ADAR1p110 expression.

Our *Adar1p150*-specific knockout contrasts with the previous model, which used an inverted PGK-Neo cassette to disrupt the isoform (**Fig 1A**). Analysis of the available published western blots from the existing *Adar1p150^-/-^* model indicates that the retained ADAR1p110 may not be expressed comparably to the *Adar1p150^+/-^* sample, nor is it induced by IFN treatment in a manner similar to the control used in this study (Pestal et al., 2015; Ward et al., 2011b). Therefore, two potential caveats of the originally developed *Adar1p150^-/-^* allele are (1) the interferon-inducible expression of ADAR1p110 may be reduced due to the disruption of the IFN promoter and, (2) the inverted pPGK-Neo cassette may impact ADAR1p110 expression from both the targeted and non-targeted allele via transcriptional interference and/or production of aberrant antisense RNAs, as has been reported for other alleles using this method of gene disruption (Matthaei *et al*., 2011; Scacheri et al., 2001; Steinman and Wang, 2011; Ward *et al*., 2011c). This may be particularly relevant when the allele is paired with other mutants as a compound heterozygote (Hubbard et al., 2022; Maurano et al., 2021; Pestal *et al*., 2015).

### ADAR1p150 specific knockout mice are embryonic lethal

Having established that our allele was a *Adar1p150-null* allele, without impacting ADAR1p110 expression, we sought to characterize its phenotype. *Adar1p150^+/-^* heterozygous mice were fertile and had no discernible phenotype. We inter-crossed *Adar1p150^+/-^* animals to determine the survival of *Adar1p150^-/-^* mice. No live *Adar1p150^-/-^* pups were identified (**Fig 2A**). Based on the embryonic lethality of the *Adar1^-/-^*mice (Hartner *et al*., 2004; Hartner *et al*., 2009; Wang *et al*., 2004), we assessed embryos at E11.5 and E12.5 and found *Adar1^+/+^, Adar1p150^+/-^* and *Adar1p150^-/-^* embryos were present at the expected Mendelian ratio (**Fig 2A**). *Adar1p150^-/-^* embryos, while present, showed evidence of developmental abnormalities and reduced viability between E11.5 and E12.5. At E11.5, three of the eight *Adar1p150^-/-^* embryos had a normal appearance whereas the other five had morphological abnormalities. At E12.5, all *Adar1p150^-/-^* embryos had abnormal features. The *Adar1p150^-/-^* embryos were fragile and had a pale color. Some abnormal embryos showed hemorrhage on the back or the head (**Fig 2B**). Similar features were previously reported for the *Adar1^-/-^* mutant, lacking both p110 and p150, with embryonic lethality at E11.0-E12.5 (Hartner *et al*., 2004; Hartner *et al*., 2009; Wang *et al*., 2004). Given the importance of ADAR1 in inhibiting the activation of MDA5 and its downstream ISGs, we performed qRT-PCR on tissue from the embryos. At both E11.5 and E12.5, *Adar1p150^-/-^* animals had a profound induction of the ISGs *Ifit1* and *Irf7* (**Fig 2C**) compared to the littermate wildtype controls. The expression of *Ifit1* and *Irf7* was higher at E12.5 than in E11.5 in the *Adar1p150^-/-^* embryos (**Supplemental Fig 1B**). The level of ISGs in the previous *Adar1p150^-/-^* mutant was not investigated. The embryonic lethality of our *Adar1p150^-/-^* is consistent with the originally described *Adar1p150^-/-^* allele, which died between E11.0-E12.0 (Ward *et al*., 2011c).

**Fig 2.**
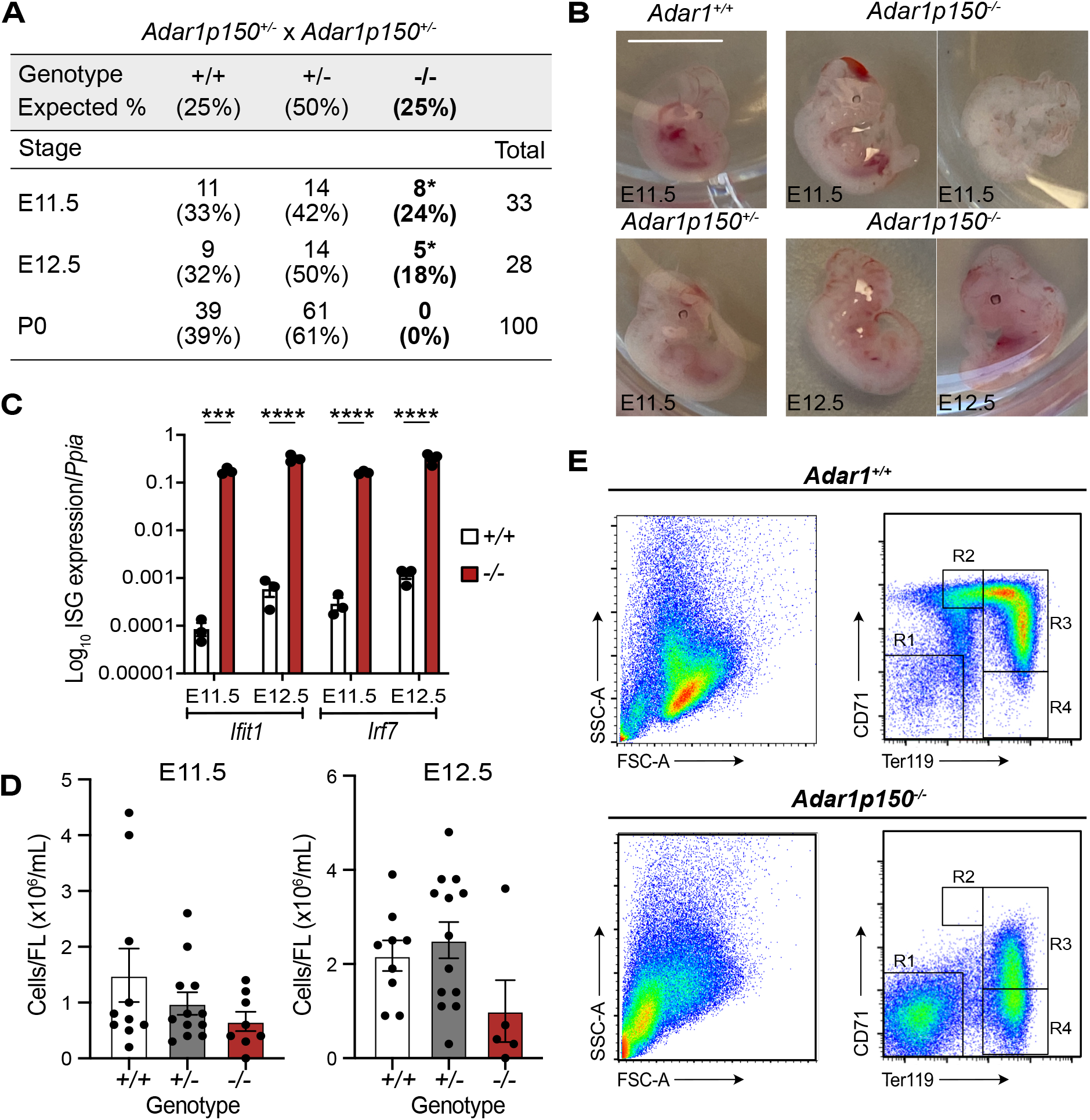
ADAR1p150 specific knockout mice die mid-gestation at E11.5-E12.5. **(A)** Survival and frequency of *Adar1^+/+^ Adar1p150^+/-^* and *Adar1p150^-/-^* mice at three development stages E11.5, E12.5 and P0 (post birth) from *Adar1p150^+/-^* inter-crosses. **(B)** Representative images of embryos of the indicated genotypes at E11.5 or E12.5. The scale bar represents 0.5 cm. **(C)** Expression of *Ifit1* and *Irf7*, both interferon-stimulated genes (ISGs), in *Adar1^+/+^* and *Adar1p150^-/--^* embryos, at E11.5 and E12.5. Data expressed as Log_10_ gene expression relative to *Ppia* expression (reference gene). Three independent samples were used for each indicated genotype and developmental stage. Significance was determined by a twoway ANOVA test with multiple comparisons with statistical significance of ***P <0.001 and ****P <0.0001. Data represent the mean +/- SEM. **(D)** Cellularity of fetal liver from E11.5 (left) and E12.5 (right). At E11.5, the number of *Adar1^+/+^* (+/+) n=10, *Adar1p150^+/-^*(+/-) n=12 and *Adar1p150^-/-^*(-/-) n=8. At E12.5, the number of +/+ n=9, +/- n=13 and -/- n=5. Significance was determined by a One-way ANOVA test with multiple comparisons. Error bars are SEM. **(E)** Representative flow cytometry profiles of viable FL erythroid cells at E12.5 with SSC-A/FSC-A (left); the proportion of erythroblast population labelled with CD71/Ter119 (right). R1=CD71^-^Ter119^-^ R2=CD71^high^Ter119^med^, R3=CD71^high^Ter119^high^ and R4=CD71^-^ Ter119^high^.

Detailed phenotypic analysis of *Adar1p150^-/-^* embryos has not been reported. Previous studies found that both *Adar1* null (*Adar1^-/-^*) and the editing deficient (*Adar1^E861A/E861A^*) embryos had a failure in hematopoiesis in fetal liver (FL)(Hartner *et al*., 2004; Hartner *et al*., 2009; Liddicoat *et al*., 2015; Wang *et al*., 2004). To determine the effects of loss of ADAR1p150 in hematopoiesis, we assessed the cells within the FL by flow cytometry. *Adar1p150^-/-^* FL had fewer viable cells at both E11.5 and E12.5, although there was no statistical difference (**Fig 2D).** In one of the *Adar1p150^-/-^* FLs that had more viable cells, the p150-deficient FL had no phenotypic erythroid cells (**Fig 2E, left**). The *Adar1p150^/-^* embryo also had a loss of CD71^high^Ter119^high^ (R3) nucleated erythroblasts and an accumulation of CD71^-^Ter119^-^ (R1) cells (**Fig 2E, right)**, indicating defective erythropoiesis in the FL in the absence of ADAR1p150.

These data demonstrate that the ADAR1p150 isoform is essential for embryonic development *in vivo*. The embryonic lethality, abnormal embryo appearance, abnormal fetal liver hematopoiesis and high ISG signature of the *Adar1p150^-/-^* embryos reproduced the previously described phenotypes observed in both *Adar1^-/-^* or *Adar1^E861A/E861A^* mutants (Hartner *et al*., 2004; Hartner *et al*., 2009; Liddicoat *et al*., 2015; Wang *et al*., 2004). Taken together, the essential physiological function of ADAR1 in fetal hematopoiesis and suppressing interferon signaling are mediated by ADAR1p150. The expression of ADAR1p110 cannot compensate for this function.

### ADAR1p150 is essential for adult homeostasis and hematopoiesis in vivo

To understand the specific requirement for ADAR1p150 in adult homeostasis we used an inducible model that we have previously used to assess the somatic restricted expression of the editing deficient ADAR1 mutation (Heraud-Farlow *et al*., 2017; Liddicoat *et al*., 2015). We crossed the *Adar1p150^+/-^* animals to *R26*-CreER^T2^ *Adar1^fl/fl^* mice. This enables tamoxifen-induced deletion of the floxed *Adar1* allele, leaving mice either heterozygous and retaining expression of p110 and p150 isoforms (*fl/+* becomes *Δ/+*) or only expressing the p150-deficient allele (*fl/p150-* become *Δ/p150-). R26*-CreER *Adar1^fl/+^* (control) and *R26*-CreER *Adar1^fl/p150-^* adult mice aged 8 to 10 weeks were fed tamoxifen-containing food for up to 28 days (**Fig 3A**). This bypassed the developmental requirement for ADAR1p150. The *Adar1* floxed allele is deleted broadly across cell types and organs upon tamoxifen treatment. We confirmed the efficient recombination of the floxed allele in the bone marrow of the *Δ/+* mice (**Fig 3B; Supplemental Fig 2A)**. In contrast, despite being moribund the *Δ/p150-* mice retained a high percentage of the *Adar1* floxed allele, indicating most likely selection against cells that had deleted the floxed allele. This phenomenon of retention of an unexcised allele has previously been seen using the same experimental model with both a *fl/fl* (loss of *Adar1* in both alleles upon tamoxifen) and *fl/E861A* (expressing editing dead ADAR1 upon tamoxifen) genotypes (Heraud-Farlow *et al*., 2017; Liddicoat *et al*., 2015).

**Fig 3.**
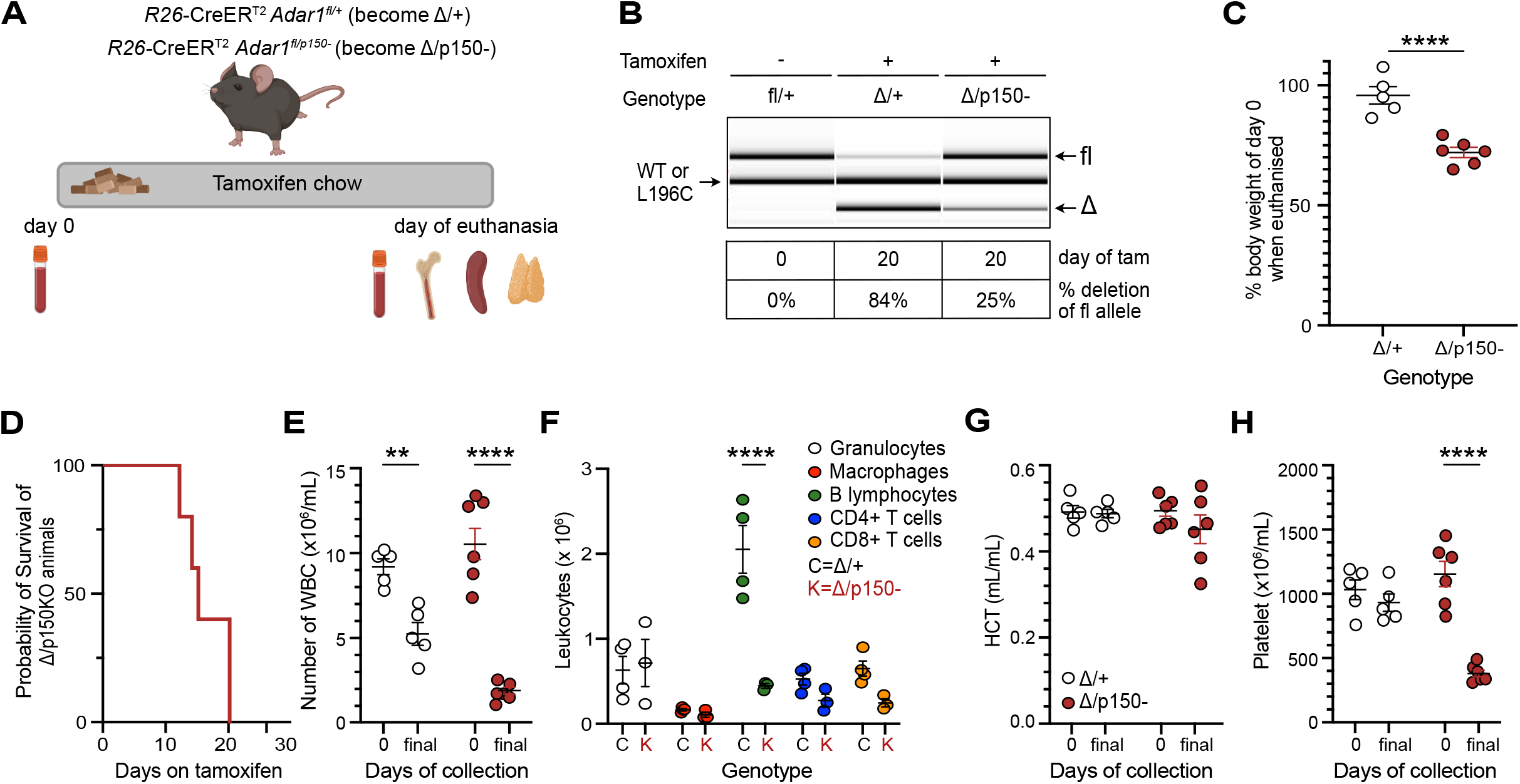
Somatic deletion of ADAR1p150 in adult mice is lethal. **(A)** Illustration of somatic deletion model. Upon tamoxifen treatment, *R26-*CreER *Adar1^fl/+^ and R26-CreER Adar1^fl/p150-^* animals became *Δ/+* and *Δ/p150-*, respectively. **(B)** Representative genotyping of genomic DNA and percentage deletion of the floxed allele on the day of euthanasia using DNA isolated from whole BM. Image and recombination percentages were calculated using LabChip (PerkinElmer). **(C)** Percentage change in body weight on the day of euthanasia compared to day 0 (before the start of the tamoxifen diet). *Δ/+* n=5 and *Δ/p150-* n =6. **(D)** Kaplan-Meier survival plot of the *Δ/p150-* animals (n=6). Note the control animals are not plotted and were healthy; each *Δ/p150-* was collected with a control animal on the same day to allow paired analysis. **(E)** Total white blood cell counts in PB at day 0 (pre-tamoxifen) and day of euthanasia (final). **(F)** Absolute numbers of each lineage in PB on the day of euthanasia (final). **(G)** Hematocrit (HCT) populations in PB at day 0 (pre-tamoxifen) and day of euthanasia (final). **(H)** Total platelets in the PB at day 0 (pre-tamoxifen) and day of euthanasia (final). Statistical comparison in all plots was done by unpaired T-test (panel I), one-way (panel E, G and H) or two-way (panel F) ANOVA tests with multiple comparisons with statistical significance of *P <0.05; **P <0.01, ***P <0.001 and ****P <0.0001. Data represented as mean +/- SEM.

The loss of ADAR1p150 led to significant weight loss compared to the day 0 weights, resulting in euthanasia prior to day 28 due to meeting of ethical endpoints (**Fig 3C**). Each moribund *Δ/p150-* animal was assessed in parallel with a paired control *Δ/+* on the same day (**Fig 3D**; note that the *Δ/+* are not represented on the plot and have survived to 28 days of tamoxifen feeding without requiring euthanasia in all prior experiments). Given the characterized impact of ADAR1 loss on hematopoiesis (Hartner *et al*., 2004; Hartner *et al*., 2009; Liddicoat et al., 2016), we quantified peripheral blood (PB) indices of mice on day 0 and on the day of collection (**Fig 3 E-H; Supplemental Fig 2B-F**). The loss of ADAR1p150 in the *Δ/p150-* mice resulted in significant declines in the total PB leukocyte populations (**Fig 3E**), mostly driven by reductions in B lymphocyte numbers (**Fig 3F**), as well as reductions in platelets (**Fig 3H**) and reduced mean corpuscular volume (**Supplemental Fig 2D**).

We further assessed the requirement for ADAR1p150 in hematopoiesis by collecting and analyzing the bone marrow (BM), spleen and thymus on the day of euthanasia (**Fig 4 Supplemental Fig 2G-I)**. In the BM, there was a reduction in the total cellularity (**Fig 4A**) and a significant loss of CD71+/Ter119+ erythroid cells (**Fig 4B**). The number of multipotent common myeloid progenitors (CMP) was significantly decreased (**Fig 4C**), and the phenotypic populations of haemopoietic stem and progenitor cells were abnormal (**Fig 4D**) (Oguro et al., 2013). The activation of the innate immune/interferon pathway was assessed by measuring the cell surface expression of Sca-1, an IFN-induced cell surface protein, by flow cytometry. There was a ~10-fold upregulation of Sca-1 expression on the BM cells of p150 deficient animals (**Fig 4E-F**). The expression of two hallmark ISGs, *Ifit1* and *Irf7*, was assessed by qRT-PCR in BM cells. There was an elevated expression of both genes in the *Δ/p150-* BM (**Fig 4G-H**). In the spleen, there was reduced weight and cellularity (**Fig 4I; Supplemental Fig 2H**). This was due to reduced numbers of B lymphocytes (**Fig 4J**) and erythroid cells (**Fig 4K**). The thymus had reduced cellularity and weight (**Fig 4L; Supplemental Fig 2I**), with the numbers of CD4+/CD8+ T cells drastically reduced (**Fig 4M**). These results demonstrate a continuous requirement for ADAR1p150 for normal homeostasis of these organs, consistent with a generalized failure in hematopoiesis as seen in both *Adar1* null and editing dead models (Hartner *et al*., 2009; Heraud-Farlow *et al*., 2017; Liddicoat *et al*., 2016; Liddicoat *et al*., 2015).

**Fig 4.**
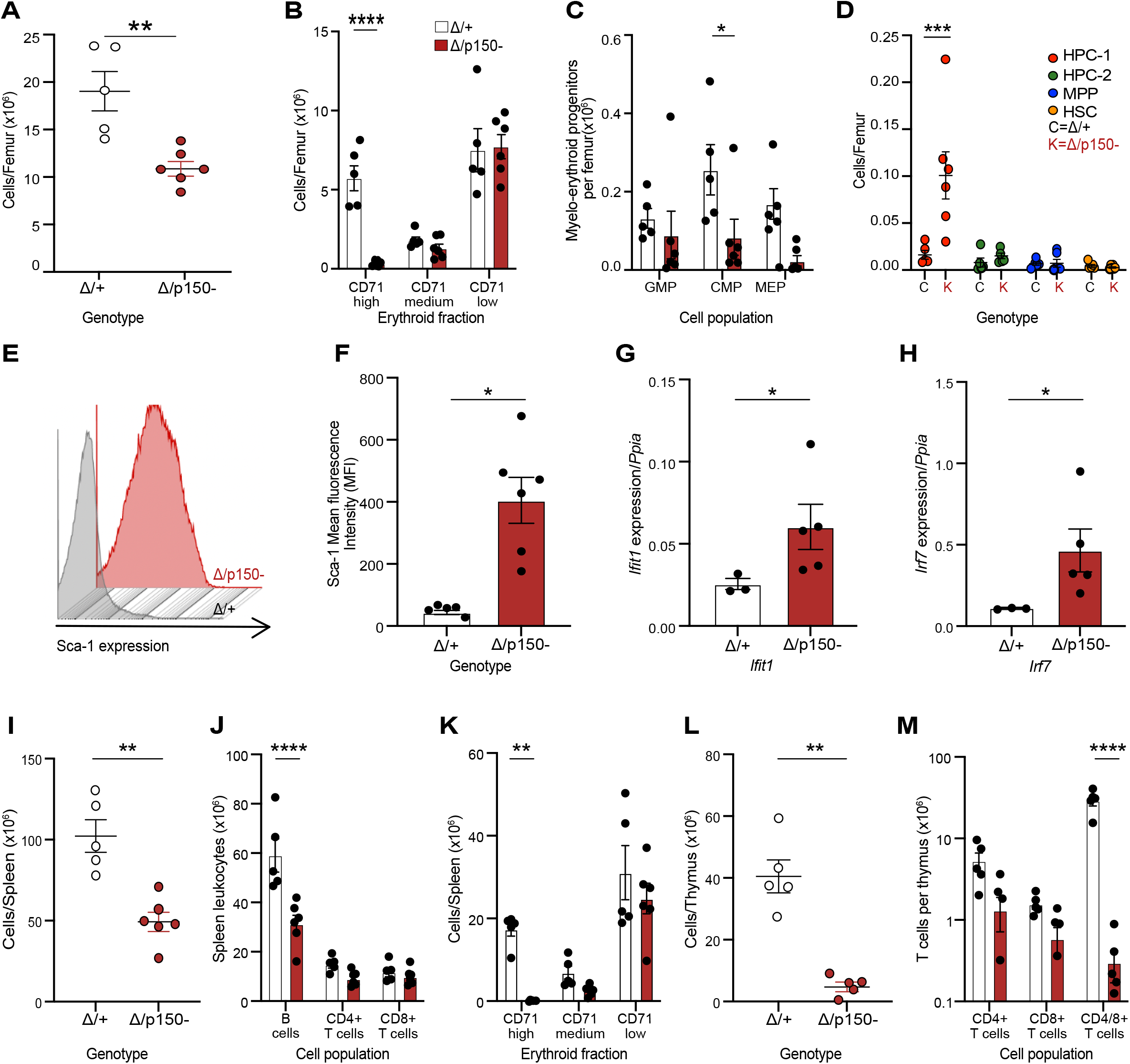
Acute loss of ADAR1p150 disrupted hematopoietic homeostasis in adult mice accompanied by an elevated innate immune response. Analysis of bone marrow, spleen and thymus from *R26-*CreER *Adar1^Δ+^ and R26-*CreER *Adar1^Δ/p150-^* animals described in Fig.3-4. **(A)** Total cellularity per femur. **(B)** Erythroid cells per femur. **(C)** Numbers of myelo-erythroid progenitors per femur. **(D)** The absolute number of stem cell and multipotent progenitor populations per femur. **(E)** Representatives flow cytometry histograms of Sca-1 expression on the lineage negative (lin^-^c-kit^+^) bone marrow fraction. **(F)** Quantification of the mean Sca-1 fluorescence intensity by flow cytometry. **(G)** Normalized expression of *Ifit1* transcripts measured by qRT-PCR in the bone marrow; *Δ/+* (n=3) and *Δ/p150-* (n=5). Data expressed as mean +/- SEM gene expression relative to *Ppia* expression. **(H)** Normalized expression of *Irf7* transcripts measured by qRT-PCR in the bone marrow; *Δ/+* (n=3) and *Δ/p150-* (n=5). Data expressed as mean +/- SEM gene expression relative to *Ppia* expression. **(I)** Total viable cellularity of the spleen. **(J)** The absolute number of B lymphocytes, CD4 positive (CD4+) and CD8 positive (CD8+) T lymphocytes in the spleen. **(K)** Erythroid cells in the spleen. **(L)** Total viable cellularity of the thymus. **(M)** The absolute number of CD4 positive (CD4+), CD8 positive (CD8+) and CD4 and CD8 double positive (CD4/8+) T cells in the thymus. Statistical tests used were unpaired T-test (panels A, F, G, H, I and L) and two-way ANOVA (panels B, C, D, J, K and M) with statistical significance of *P <0.05; **P <0.01, ***P <0.001, ****P <0.0001. Data expressed as mean +/- SEM. In panels A-F and I-M *Δ/+* n=5 and *Δ/p150^-^* n=6.

The new ADAR1p150-specific knockout mouse model described here has allowed characterization of the *in vivo* functions of ADAR1p150. We demonstrate that this allele resulted in the specific loss of the ADAR1p150 isoform, without any apparent impacts on the expression of the retained p110 isoform, both basally and following IFN β treatment. We completed analysis that allows a direct comparison to previous studies describing the phenotypes of *Adar1* null (loss of both p110 and p150) and editing deficient mutants (E861A)(Hartner *et al*., 2004; Hartner *et al*., 2009; Liddicoat *et al*., 2015; Wang *et al*., 2004). This provides an important comparison of these distinct alleles. When considered together with the recently described *Adar1p110^-/-^* allele, we conclude that ADAR1p150 is specifically required to suppress the activation of the innate immune response and allow normal development and adult homeostasis. This is the physiologically essential and non-redundant function of ADAR1p150.

## Materials and Methods

### Ethics Statement

All animal experiments were approved by the Animal Ethics Committee of St. Vincent’s Hospital, Melbourne, Australia (Protocol number #016/20). Animals were euthanized by CO2 asphyxiation or cervical dislocation.

### Animals

*Adar1^L196C^* mice were identified as an incidental mutation arising from CRISPR/Cas9 targeting in C57BL/6 zygotes to generate a p.P195A knock-in point mutation by the Monash Genome Modification Platform (Monash University, Clayton, Australia)(Liang *et al*., 2022). A single nucleotide deletion at nucleotide 587 (587delT) resulted in the p.L196CfsX6 mutation (nomenclature based on NCBI CCDS50963). The introduction of the mutation was confirmed by Sanger sequencing of the region in both the founders and subsequent generations. *Adar^fl/fl^* (*Adar1^fl/fl^;* exon 7-9 floxed; MGI allele: *Ada^tm1.Phs^;* MGI:3828307) (Hartner *et al*., 2004; Hartner *et al*., 2009) and *Rosa26*-CreER^T2^ (Gt(ROSA)26Sor^tm1(cre/ERT2)T^yj) (Ventura et al., 2007) mice were on a backcrossed C57BL/6 background as previously described (Hartner *et al*., 2009; Heraud-Farlow *et al*., 2017; Liddicoat *et al*., 2016; Liddicoat *et al*., 2015). All animals were housed at the BioResource’s Centre at St. Vincent’s Hospital, Melbourne, Australia. Mice were maintained and bred under specific pathogen-free conditions with food and water provided *ad libitum*. For the somatic deletion models *R26-*CreER *Adar1^fl/+^* and *R26*-CreER *Adar1^fl/p150^*, all animals were aged from 8 to 10 weeks at the initiation of tamoxifen; Tamoxifen containing food were prepared at 400mg/kg tamoxifen citrate (Selleckchem) in standard mouse chow (Specialty Feeds, Western Australia, Australia).

### Genotyping

Genotyping of the L196C mutants was determined by PCR and Sanger sequencing. The following primers: Primer P1 (5’-ACCATGGAGAGGTGCTGACG-3’) and P2 (5’-ACATCTCGGGCCTTGGTGAG-3’) were used to obtain a 489bp product. The purified PCR product was sequenced using P1 primer (Australian Genome Research Facility, Melbourne). Genotyping of all other lines and Cre recombination was performed as previously described by (Heraud-Farlow *et al*., 2017; Liddicoat *et al*., 2015).

### Embryo and fetal liver analysis

Timed matings of *Adar1p150^+/-^* females was undertaken for embryo analysis, and embryos were collected at E11.5 and E12.5. Fetal livers (FL) were isolated from the embryos and suspended in 1mL of PBS containing 2% FBS using a 21G needle/1mL syringe. FL cell counts were performed on a hematological analyzer (Sysmex KX1000). Single-cell FL suspensions were then subjected to antibody staining for flow cytometry analysis. The measurement of erythroid cells used antibodies against murine Ter119 (PE), CD71 (APC), and CD44 (PE-Cy7) sourced from eBioscience, BioLegend or BD Pharmingen. Cells were acquired on a BD LSRIIFortessa and analysed with FlowJo software Version 9 or 10.0 (Treestar).

### Mouse Embryonic Fibroblasts and immortalization

Mouse embryonic fibroblasts (MEFs) were generated from E11.5 embryos of the indicated genotypes. The embryos were dissected, the head was used for qRT-PCR, and the heart and fetal liver (for flow cytometry) were removed. MEFs were made from the remaining embryo. The tissue was drawn through an 18G needle/1mL syringe, suspended in 1mL of 0.025% Trypsin-EDTA (Gibco/Thermo Fisher) and incubated at 37°C in a 10cm^2^ tissue culture plate for 30 minutes. Then, 10ml of normal growth media (High glucose DMEM (Sigma) containing10% FBS (not heat-inactivated, Assay Matrix), 1% Penicillin/Streptomycin (Gibco), 1% Glutamax (Gibco) and 1% amphotericin B (Sigma; 250ug/mL stock)) was directly added to the plate. The digested tissue was re-suspended and dispersed. The MEFs were incubated in a hypoxia chamber flushed with 5% oxygen and 5% carbon dioxide in nitrogen at 37°C. Once the cells were confluent (>70% confluency), the cells were trypsinized and passaged onto 10cm^2^ plates in normoxic conditions for all further culturing. MEFs were immortalized with 1ml of media containing an shRNA targeting murine p53 (shp53.1224 in LMP vector), 1% polybrene, and normal growth media (Dickins et al., 2005). After 12 hours of infection, 1ml of fresh media was added to the cells. After 72 hours, the media was replaced with fresh normal growth media. The immortalized MEFs were treated with recombinant murine interferon beta (PBL Assay Science; PBL-12405) at 250U/mL for 24 hours in normal growth media.

### Western blot analysis

MEFs were collected by trypsinization and pellets washed in cold PBS and resuspended in RIPA buffer (20mM Tris·HCl, pH8.0, 150mM NaCl, 1mM EDTA, 1% sodium deoxycholate, 1% Triton X-100, 0.1% SDS) supplemented with 1x HALT protease inhibitor and 1x PhosSTOP phosphatase inhibitor (ThermoFisher Scientific). Lysates were used for western blot analysis as described below. Protein was quantified using the Pierce BCA protein assay kit (ThermoFisher Scientific) on an Enspire multimode plate reader (PerkinElmer). 20μg of protein extract per sample was loaded on pre-cast NuPAGE™ 10% or 4-12%, Bis-Tris polyacrylamide gels (Invitrogen) and transferred onto Immobilon-P PVDF membranes (Merck Millipore). Membranes were blocked with 5% milk in Tris-buffered saline with tween (TBST) and incubated at 4°C overnight with rat monoclonal anti-mouse ADAR1 antibody (clone RD4B11) (Liddicoat *et al*., 2015), rabbit anti-MDA-5 (Cell Signaling, D74E4) and mouse anti-Actin (Sigma Aldrich, A1978). Membranes were then probed with HRP-conjugated goat anti-rat (ThermoFisher Scientific, 31470), anti-rabbit (ThermoFisher Scientific, 31460) or anti-mouse (ThermoFisher Scientific, 31444) secondary antibodies and visualized using ECL Prime Reagent for chemiluminescent detection on Hyperfilm ECL (Amersham) or iBright FL1500 Imaging system (ThermoFisher). Western band intensities were quantified using Fiji.

### Peripheral blood analysis

Peripheral blood (PB) was obtained via retro-orbital bleeding into BDMicrotainer K2E tubes; (Becton Dickinson) from the somatic mutation models. PB samples were counted on a hematological analyser (Sysmex KX1000). The red blood cells were lysed using hypotonic lysis buffer (150mM NH4Cl, 10mM KHCO3, 0.1mM Na2EDTA, pH7.3) and resuspended in 50μl of FACS buffer for flow cytometry analysis.

### Flow cytometry analysis

Antibodies against murine B220 (APC-eFluor780), CD11b/Mac-1 (PE), Gr1 (PE-Cy7), F4/80 (APC), CD4 (eFluor450) and CD8a (PerCP-Cy5.5), Ter119 (PE), CD71 (APC), CD44 (PE-Cy7), Sca-1(PerCP-Cy5.5), c-Kit (APC-eFluor780), CD150 (PE), CD48 (PE-Cy7), CD34(eFluor660), CD16/32 (eFluor450) and biotinylated antibodies (CD2, CD3e, CD4, CD5, CD8a, B220, Gr-1, CD11b/Mac1) were used. The biotinylated antibodies were detected with streptavidin-conjugated Brilliant Violet 786. All antibodies were obtained from eBioscience, BioLegend or BD Pharmingen. (Heraud-Farlow *et al*., 2017; Liddicoat *et al*., 2015; Singbrant et al., 2011; Smeets et al., 2014). Cells were acquired on a BD LSRIIFortessa and analysed with FlowJo software Version 9 or 10.0 (Treestar).

### qRT-PCR

Whole heads from embryos or single cell suspensions of bone marrow from the somatic deletion models were collected and immediately snap frozen in liquid nitrogen or dry ice and stored at −80 °C. Frozen tissues were homogenised in Trisure reagent (Bioline) using IKA T10 basic S5 Ultra-turrax Disperser. RNA was isolated using Direct-Zol columns (Zymo Research) following the manufacturer’s instructions and cleaned up with Zymo clean and concentrate kit. Complementary DNA (cDNA) was synthesised using the Tetro cDNA synthesis kit (Bioline). qRT-PCR of ISGs (*Ifit1 and Irf7*) was performed using SYBR green and the ΔΔCT method (normalised to *Ppia*) as previously described (Heraud-Farlow *et al*., 2017). Duplicate or triplicate reactions per sample were measured using an AriaMx Real-time PCR machine (Agilent). The following primers were used: *Ifit1* P1 (5’-ATGGGAGAGAATGCTGATGG −3’); *Ifit1* P2 (5’-AGGAACTGGACCTGCTCTGA-3’); *Irf7* P1 (5’-CCAGTTGATCCGCATAAGGT-3’); *Irf7* P2 (5’-AGCATTGCTGAGGCTCACTT-3’); *Ppia* P1 (5’-GTCAACCCCACCGTGTTCTT −3’) and *Ppia* P2 (5’-CTGCTGTCTTTGGAACTTTG-3’).

### Statistical analysis and figure preparation

To determine statistical significance, unpaired T-tests, ordinary one-way or two-way ANOVA tests with multiple comparisons were conducted in GraphPad Prism software version 9 (GraphPad). Throughout this study, significance is indicated using the following convention: *P<0.05; **P<0.01; ***P<0.001; ****P<0.0001. Data are presented as mean ± S.E.M. The number of samples used is described in the corresponding figure legends. The figures were generated using BioRender.com and Affinity Designer.

## Acknowledgements

The authors thank S Taylor, A Goradia and E Tonkin for technical assistance; R Dickins (Australian Centre for Blood Disease, Monash University) for p53.1224 shRNA; the Monash Genome Modification Platform (MGMP) at Monash University for the generation of the *Adar1^L196C^* (p.L196C*fs*X6) mice; Monash Antibody Technology Facility (MATF) for purification of ADAR1 antibody from hybridomas; and St.Vincent’s Hospital BioResource’s Centre for the care of experimental animals. The *Adar1^L196C^* mutant mice were produced via CRISPR/Cas9 mediated genome editing by the Monash Genome Modification Platform (MGMP), Monash University as a node of Phenomics Australia. Phenomics Australia is supported by the Australian Government Department of Education through the National Collaborative Research Infrastructure Strategy, the Super Science Initiative, and the Collaborative Research Infrastructure Scheme. This work was supported by the National Health and Medical Research Council (NHMRC; APP1183553 to C.R.W and J.H.F; APP1182453 to J.H.F); a Melbourne Research Scholarship (to Z.L. from The University of Melbourne); J.H.F is supported by a fellowship from 5point Foundation; and in part by the Victorian State Government Operational Infrastructure Support Scheme to St Vincent’s Institute.

## Author contribution statement

J.H-F and C.R.W conceptualised the study. Z.L, J.H-F and C.R.W designed the experiments. Z.L, J.H-F and C.R.W performed the experiments. Z.L, J.H-F and C.R.W wrote the manuscript, and all authors reviewed and edited the manuscript; J.H-F and C.R.W were responsible for funding acquisition. J.H.F and C.R.W provided supervision.

## Supplemental Material

**Supplemental Fig 1.**
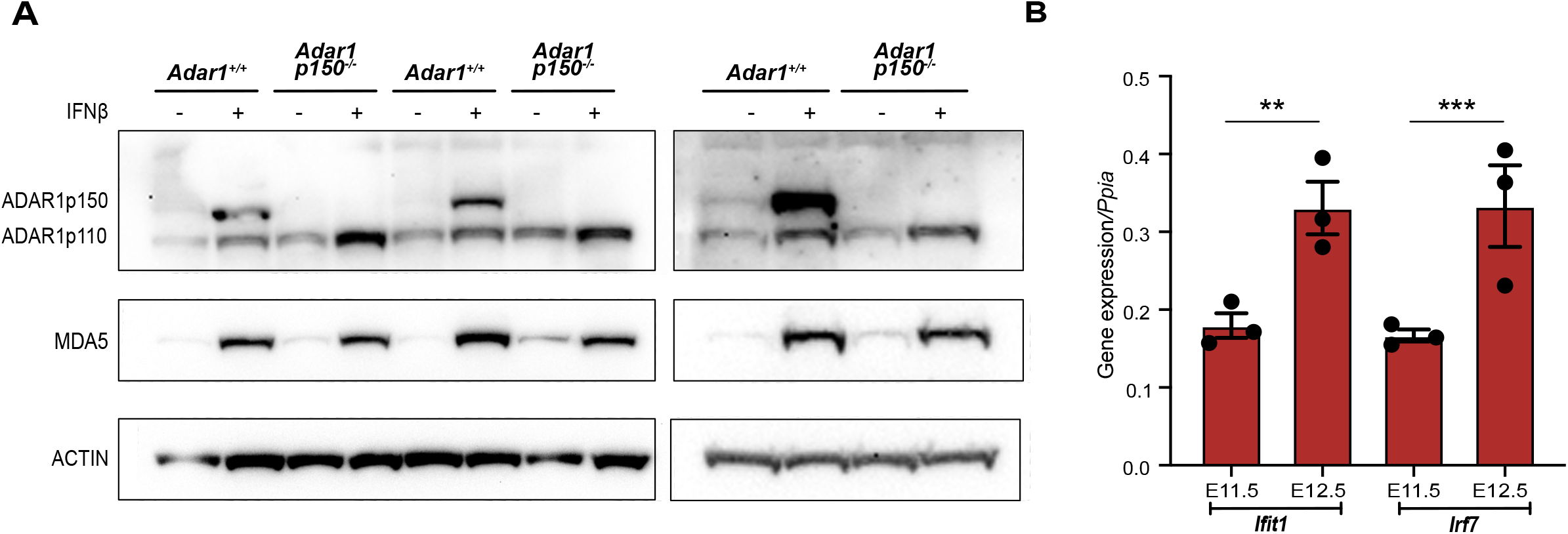
The analysis of MEFs and embryos. **(A)** Western blot analysis of *Adar1^+/+^* and *Adar1p150^-/-^* MEFs generated from 3 independent mice of each genotype with and without 24 hours IFNβ treatment. The relative expression of p110 is presented in Fig 1F. **(B)** Expression of the interferon-stimulated genes (ISGs) *Ifit1* and *Irf7* in the heads of *Adar1p150^-/-^* embryos at E11.5 and E12.5. Data expressed as mean +/- SEM gene expression relative to *Ppia* expression. Three independent embryo samples were used for each indicated genotype. Significance was determined by a two-way ANOVA test with multiple comparisons with statistical significance of ***P <0.001 and ****P <0.0001. Error bars are SEM.

**Supplemental Fig 2.**
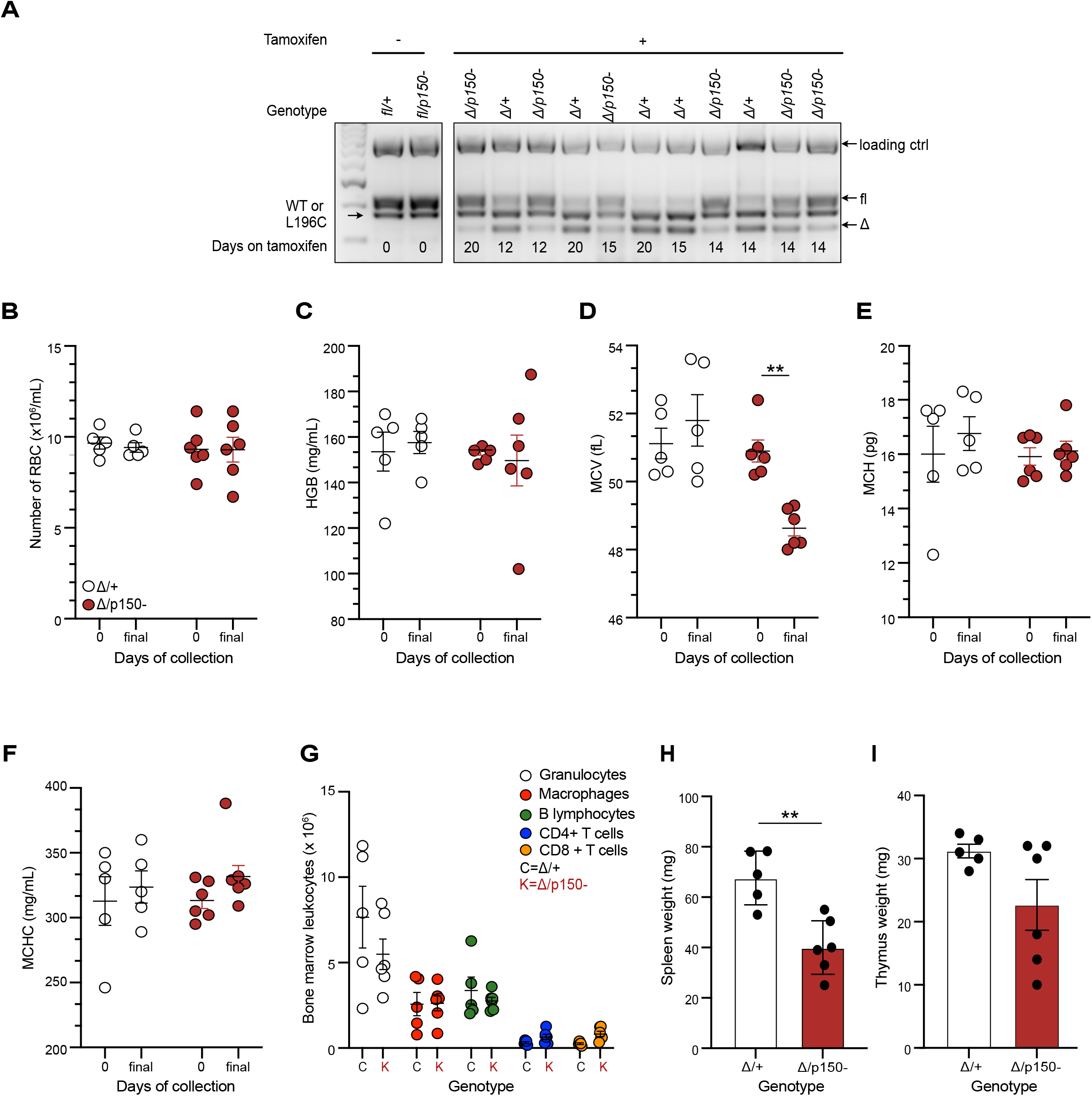
Cell counts and hematopoietic stem cell populations in PB, BM, spleen and thymus of the somatic mouse models. **(A)** Genotyping of genomic DNA from all animals used in the somatic deletion model on the day of euthanasia using DNA isolated from whole BM. **(B)** Total red blood cell counts in PB at day 0 (pre-tamoxifen) and day of euthanasia (final). **(C)** Haemoglobin level in PB at day 0 (pre-tamoxifen) and day of euthanasia. **(D)** Mean corpuscular volume of PB at day 0 (pre-tamoxifen) and day of euthanasia. **(E)** Mean corpuscular haemoglobin level in PB at day 0 (pre-tamoxifen) and day of euthanasia (final). **(F)** The mean corpuscular haemoglobin concentration in PB at day 0 (pre-tamoxifen) and day of euthanasia(final). **(G)** Differential analysis of leukocyte populations in the BM on day of euthanasia. **(H)** Weights of the spleen. **(I)** Weights of the thymus. Statistical tests used were two-way ANOVA (B-G) and unpaired T-test (H-I) with statistical significance of *P <0.05; **P <0.01, ***P <0.001, ****P <0.0001. Displayed as mean +/- SEM. *Δ/+* n= 5, *Δ/p150-* n=6.

